# Validation and Establishment of a SARS-CoV-2 Lentivirus Surrogate Neutralization Assay as a pre-screening tool for the Plaque Reduction Neutralization Test

**DOI:** 10.1101/2022.09.13.507876

**Authors:** John Merluza, Johnny Ung, Kai Makowski, Alyssia Robinson, Kathy Manguiat, Nicole Mueller, Jonathan Audet, Julie Chih-Yu Chen, James E Strong, Heidi Wood, Alexander Bello

## Abstract

Neutralization assays are important in understanding and quantifying neutralizing antibody responses towards SARS-CoV-2. The SARS-CoV-2 Lentivirus Surrogate Neutralization Assay (SCLSNA) can be used in biosafety level 2 (BSL-2) laboratories and has been shown to be a reliable, alternative approach to the plaque reduction neutralization test (PRNT). In this study, we optimized and validated the SCLSNA to assess its ability as a comparator and pre-screening method to support the PRNT. Comparability between the PRNT and SCLSNA was determined through clinical sensitivity and specificity evaluations. Clinical sensitivity and specificity produced acceptable results with 100% (95% CI: 94-100) specificity and 100% (95% CI: 94-100) sensitivity against ancestral Wuhan spike pseudotyped lentivirus. The sensitivity and specificity against B.1.1.7 spike pseudotyped lentivirus resulted in 88.3% (95% CI: 77.8 to 94.2) and 100% (95% CI: 94-100), respectively. Assay precision measuring intra-assay variability produced acceptable results for High (1:≥ 640 PRNT_50_), Mid (1:160 PRNT_50_) and Low (1:40 PRNT_50_) antibody titer concentration ranges based on the PRNT_50_, with %CV of 14.21, 12.47, and 13.28 respectively. Intermediate precision indicated acceptable ranges for the High and Mid concentrations, with %CV of 15.52 and 16.09, respectively. However, the Low concentration did not meet the acceptance criteria with a %CV of 26.42. Acceptable ranges were found in the robustness evaluation for both intra-assay and inter-assay variability. In summary, the validation parameters tested met the acceptance criteria, making the SCLSNA method fit for its intended purpose, which can be used to support the PRNT.

## Introduction

The COVID-19 pandemic has caused an unprecedented amount of 448,624,192 confirmed cases and 6,507,879 deaths worldwide as of September 9, 2022 (https://www.worldometers.info/coronavirus/). However, the rapid development and administration of vaccines such as Pfizer-BioNTech and Moderna have contributed in helping prevent severe disease and mortality among infected individuals (1–3). As the COVID-19 pandemic unfolded over time, it was shown that the spike glycoprotein found in SARS-CoV-2 virus membrane can undergo mutations resulting in variants that can evade neutralizing antibodies generated against previous iterations of spike, leading to new waves of infection (4, 5). Breakthrough infections have been a challenge throughout the pandemic and neutralization studies are important in analyzing the neutralizing antibody response, which plays an essential role in preventing severe infection and for assessing vaccine candidate suitability (6, 7).

The PRNT assay is the current gold-standard neutralization assay; however, this method is labor intensive and requires the use of a Biosafety Level 3 (BSL-3) or higher containment laboratory (8–12). In addition, the PRNT assay relies on visualization of plaques formed by the virus, resulting in longer turnaround time (TAT) from sample receipt to result (13, 14). Such limitations present challenges in sample processing and throughput capabilities and alternate methodologies are required to help circumvent these difficulties. The SCLSNA is one such approach that does not have the same logistical challenges associated with the PRNT assay; SCLSNA can be safely performed in BSL-2 laboratories, it is amenable to high-throughput and has a relatively faster TAT of 48 hours (15–18). The SCLSNA incorporates the use of lentiviruses pseudotyped with SARS-CoV-2 spike protein, which can serve as a surrogate virus to quantitate neutralizing antibodies generated against the SARS-CoV-2 spike protein (19, 20). The lentivirus particles used in this study are second generation lentiviral vectors that do not contain accessory virulence genes such as *vif, vpu* and *nef*, rendering them replication incompetent and allowing for safe use in a BSL-2 laboratory (21).

In this study, we performed a method validation to determine if the SCLSNA is fit for its intended purpose as a reliable comparator and screening method to complement the PRNT (22). Following guidelines recommended by the WHO and Food and Drug Administration (FDA), this study targeted validation parameters such as precision, repeatability, robustness, linearity, LOD and LOQ (22, 23). We optimized the SCLSNA to confirm optimal assay parameter conditions and to limit variation, as well as assess clinical sensitivity and specificity studies in comparison to the PRNT.

## Materials and Methods

### Study population and specimen collection

Plasma samples used in the validation were obtained from the National Microbiology Laboratory NML COVID-19 National Panel (NML CNP) under the approval of the Research Ethics Board (REB-2020-004P). Samples from patients that previously tested positive for SARS-CoV-2 by quantitative reverse transcription PCR (RT-qPCR) were included in the NML COVID-19 National Panel. Blood draw collection dates took place from May 13, 2020 to August 22, 2020. All samples were collected from various provinces nation-wide through the Canadian Blood Services (24). Plasma samples were heat-inactivated for 30 minutes at 56°C, then stored at -80°C until testing was performed.

### Cell lines

For the SCLSNA, HEK293T/ACE2-TMPRSS2 cells (GeneCopoeia™, Rockville, MD) were used for infection by pseudotyped lentivirus. These cells stably express angiotensin-converting enzyme 2 (ACE2) and transmembrane serine protease 2 (TMPRSS2), which are important for infection by SARS-CoV-2 and other pseudotyped viruses expressing SARS-CoV-2 spike on their surface.

For pseudotyped lentivirus production, AAVpro 293T cells (TaKaRa Bio, San Jose, CA) were used to transfect the envelope plasmid, transfer plasmid and packaging plasmid. All cell lines were incubated in a 5% CO_2_ incubator at 37°C with DMEM (Gibco, Waltham, MA) supplemented with 10% heat-inactivated fetal bovine serum, 1% penicillin/streptomycin, 1% L-glutamine and 1% sodium pyruvate (DMEM10) (Gibco, Waltham, MA).

### SARS-CoV-2 pseudotyped virus production

All assays and lentivirus preparations were performed in BSL-2 conditions unless noted differently. AAVpro 293T cells (Takara Bio, San Jose, CA) were used to generate the SARS-CoV-2 spike pseudotyped lentiviruses in 10 – 150 mm Corning dishes (Millipore Sigma, St.Louis, MO). Briefly, psPAX2 empty vector HIV packaging plasmid (addgene, Watertown, MA, a gift from Didier Trono), SARS-CoV-2 spike protein (ancestral Wuhan or B.1.1.7) envelope expression plasmid (GeneCopoeia™, Rockville, MD) and pHAGE-CMV-Luc2-IRES-ZsGreen-W transfer vector plasmid (a kind gift from Jesse Bloom) were transfected into each plate at 42.19 μg, 60.94 μg, and 60.94 μg, respectively. Transfections were performed using CalPhos Mammalian Transfection Kit (TaKaRa Bio, San Jose, CA) and plates incubated in a 5% CO_2_ incubator at 33°C for 16 hours with DMEM10. Following incubation, the media was replaced with 11 mL of fresh DMEM10 and incubated for an additional 18-24 hours.

Culture supernatants were clarified by centrifugation at 500g, 4°C for 5 minutes using a Sorvall ST-40R, TX-1000. Supernatants were pooled and filtered using a 0.45 μm PES filter (ThermoFisher Scientific, Waltham, MA) and ultracentrifuged in ultra-clear round bottom tubes (FisherScientific, Waltham, MA) using an Optima™ L-90K ultracentrifuge and SW 32 Ti Swinging-Bucket Rotor at 16°C for 2.5 hours at 50 000g. Pellets were resuspended in 1X PBS, aliquoted and stored at -80°C.

### SARS-CoV-2 Lentivirus Surrogate Neutralization Assay

Neutralization was measured by the reduction of luciferase expression for samples incubated with pseudotyped lentivirus relative to luciferase expression in control wells containing only SARS-CoV-2 pseudotyped lentivirus and cells. The half-maximal inhibitory concentration (IC_50_) was used as the reportable value for the SCLSNA and generated using GraphPad Prism software v.9.3. Sample dilutions were logarithm transformed (log10) and all raw data were normalized to a common scale(25). For data normalization, the wells containing pseudotyped lentivirus + HEK293T/ACE2-TMPRSS2 cells were defined as “0% neutralization” and wells containing only HEK293T/ACE2-TMPRSS2 cells were defined as “100% neutralization”. A nonlinear regression curve was used to determine the IC_50_ values for the samples once the relative luminescence units (RLU) decreased to half the response of the virus control wells.

In preparation for the SCLSNA, HEK293T/ACE2-TMPRSS2 cells were seeded at 1×10^4^ cells/mL in poly-L-lysine pre-coated plates (Corning, Glendale, ARI). Cells were incubated in a 5% CO_2_ incubator at 37°C for 18-24 hours prior to performing the assay. Test samples were diluted 1:20 followed by an eight step 2-fold serial dilution. After addition of pseudotyped lentiviruses, plates containing serially diluted test sample and pseudovirus were incubated for 1 hour at 37°C. Prior to HEK293T/ACE2-TMPRSS2 cell infection, DMEM10 cell culture media was replaced with DMEM containing 5% FBS, 5 μg/mL polybrene. The diluted test sample containing pseudovirus mixture was transferred to the cell plate and incubated in 5% CO_2_ at 37°C for 48 hours. Luminescence was detected using the Bright-Glo™ Luciferase Assay System (Promega, Madison, WI) and a Biotek Cytation 1 imaging reader. Raw data was obtained on a Biotek Gen5™ Microplate reader and data analysis was performed on GraphPad Prism version v.9.3.

### SARS-CoV-2 Plaque-Reduction Neutralization Test

The SARS-CoV-2 PRNT was adapted from a previously described method for SARS-CoV-1 (26). Briefly, serially diluted serological specimens were mixed with diluted SARS-CoV-2 at 100 plaque-forming units (PFU)/100 μL in a 96-well plate. The antibody-virus mixture was added in duplicate to 12-well plates containing pre-plated Vero E6 cells. All plates were incubated at 37°C with 5% CO_2_ for 1 hour of adsorption, followed by the addition of a liquid overlay. The liquid overlay was removed after a 3-day incubation and cells were fixed with 10% neutral-buffered formalin. The monolayer in each well was stained with 0.5% crystal violet (w/v) and the average number of plaques was counted for each dilution. The reciprocal of the highest dilution resulting in at least 50% and 90% reduction in plaques (when compared with controls) were defined as the PRNT_50_ and PRNT_90_ titers, respectively. PRNT_50_ titers and PRNT_90_ titers ≥ 20 were considered positive for SARS-CoV-2 neutralizing antibodies, whereas titers < 20 were considered negative (8).

### Statistical analysis and visualization

Neutralization was determined by IC_50_ once plasma samples reduced the RLU by 50% relative to the virus control wells. Plasma sample dilutions were log-transformed, normalized and plotted using nonlinear regression to obtain the IC_50_ values. Based on the FDA guidelines, sample suitability acceptance criteria was set at 20% coefficient of variation (CV) between sample replicates and a goodness of fit (R^2^) of ≥ 0.700 (22). Assay suitability acceptance criteria within the virus and cell control replicates for each assay was set at 30% CV and a difference of ≥ 1000X (at least 3 logs above background) between the virus control and cell control RLU (19). All data analysis was performed using GraphPad Prism v.9.3 software.

Clinical specificity and sensitivity of the SCLSNA was compared to the gold-standard PRNT assay. Sixty samples from the NML COVID-19 National Panel that tested positive for SARS-CoV-2 neutralizing antibodies (NAbs) and sixty pre-COVID-19 samples negative for SARS-CoV-2 NAbs were used in the comparison. Contingency tables were generated to calculate the sensitivity and specificity values.

Repeatability (intra-assay precision) was examined to measure the degree of agreement between results from different assays of the same homogenous sample material (23). Three different concentrations were used and classified as “High” (1:≥ 640 PRNT_50_), “Mid” (1:160 PRNT_50_) and “Low” (1:40 PRNT_50_) based on our in-house PRNT_50_ titer results. Each sample was processed in triplicate on two separate weeks for a total of six determinations each. The analysts performed the assay using the same equipment and test conditions each week under approximately the same timeframe.

Reproducibility (inter-assay variability) examined the degree of agreement between individual results using the same homogenous sample material from different analysts. Three different concentrations were assessed on different days between different analysts.

Robustness was determined by examining the ability of the SCLSNA to provide analytical results of acceptable accuracy and precision under different conditions. Three concentrations of test sample were tested in triplicate on two different weeks for a total of six determinations for each sample. The detection method compared two luminometers in different operational conditions. The Agilent BioTek Cytation1 and Promega’s Glo-Max® Navigator were used for comparison.

Assay performance and acceptance criteria were based off the %CV, which measures relative variability. The acceptance range used throughout the validation for the repeatability, reproducibility and robustness was ≤ 20% following FDA guidelines (22).

To determine linearity within the SCLSNA, a WHO international reference panel for anti-SARS-CoV-2 immunoglobulin was used to assess the ability of the assay to produce results that are directly proportional to the concentration of an analyte. The reference panel (NIBSC) consisted of pooled plasma from individuals from the United Kingdom or Norway who recovered from COVID-19 (https://www.nibsc.org/documents/ifu/20-268.pdf). The negative control consists of pre-COVID-19 plasma from healthy blood donors, collected before 2019.

## Results

### Cell seeding optimization

A cell seeding optimization experiment for HEK293T/ACE2-TMPRSS2 cells was performed to determine the optimal sensitivity for the SCLSNA while trying to maximize pseudotyped lentivirus infection and minimize variation between sample replicates. A high titer sample (≥ 1:640 PRNT_50_) was tested against an ancestral Wuhan spike pseudotyped lentivirus with nine cell seeding densities ranging from 7.8×10^2^ to 2.0×10^5^ cells/well (Figure 1). The selection for the optimal cell density was based on the combination of the cell density (> 1×10^3^ cells/well), IC_50_ (> 640) and goodness of fit (R^2^ > 0.9). The results indicate IC_50_ values > 640 and R^2^ > 0.9 for cell densities between 7.8×10^2^ to 6.3×10^3^, but these seeding densities were not selected due to the potential of increased variability in SCLSNA testing observed in the lower cell densities (9). The cell densities above 1×10^4^ cells/well demonstrated a reduced IC_50_ or R^2^. Thus, the cell density of 1.0×10^4^ cells/well was selected as an optimal cell seeding density for the SCLSNA (Table 1).

**Figure 1.**
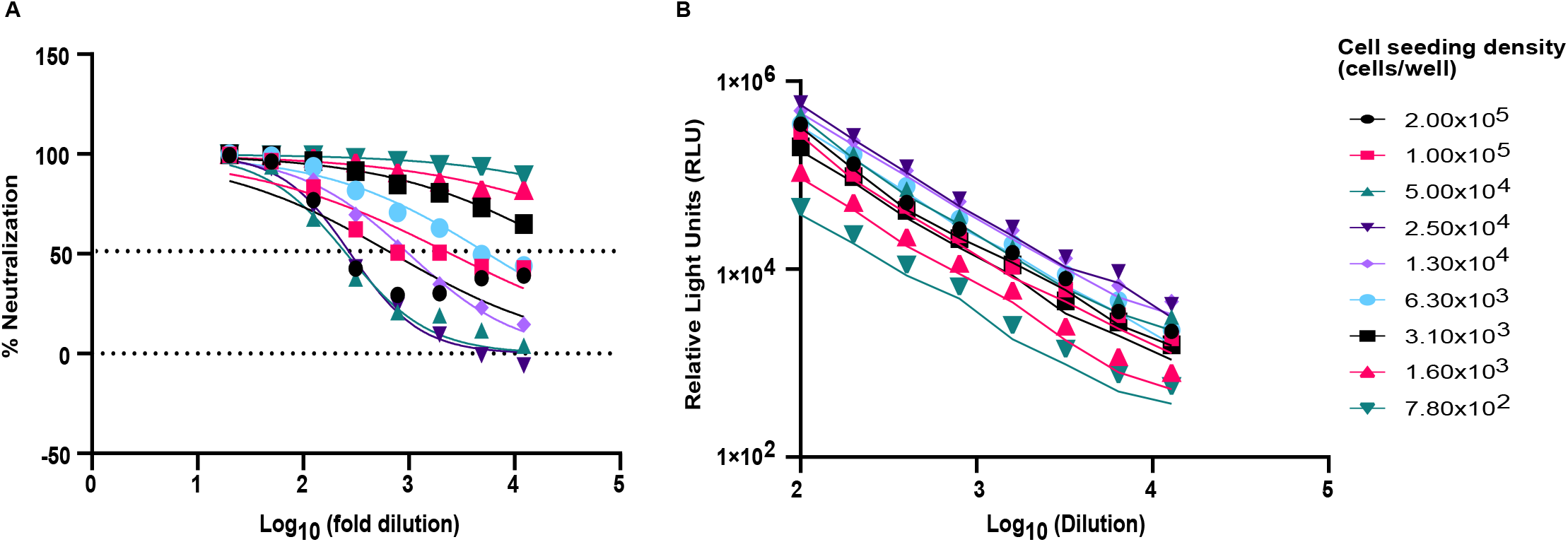
**(A)** Cell density optimization for neutralization. Cell density experiments were performed to determine optimal cell numbers based on neutralization. Nine different cell concentrations were used and performed on three plates. The average RLU values were determined for each cell concentration and used to calculate IC_50_ and R^2^ on GraphPad Prism v.9.3 software. **(B)** Pseudovirus titration against different cell seeding densities to determine optimal pseudovirus signal in relation to the cell seeding density. Ancestral Wuhan spike pseudotyped lentivirus was diluted 100-fold followed by an 8-step 2-fold serial dilution. RLU values were used to determine optimal cell seeding density. HEK293T/ACE2-TMPRSS2 cells and lentivirus were incubated in a 5% CO_2_ incubator at 37°C for 18-24 hours prior to detection of RLU.

**Table 1.** Optimal IC_50_ determination based on cell seeding density and R2 value. GraphPad Prism version 9.3 was used to determine IC_50_ and goodness of fit (R^2^) values.

| Cell density (cells/well) | IC<sub>50</sub> | R<sup>2</sup> |
| --- | --- | --- |
| 2.00x10<sup>5</sup> | 703.4 | 0.7144 |
| 1.00x10<sup>5</sup> | 2472 | 0.8515 |
| 5.00x10<sup>4</sup> | 256.8 | 0.9705 |
| 2.50x10<sup>4</sup> | 280.7 | 0.9929 |
| 1.30x10<sup>4</sup> | 997.8 | 0.9917 |
| 6.30x10<sup>3</sup> | 5.58 x10<sup>3</sup> | 0.9658 |
| 3.10x10<sup>3</sup> | 3.70 x10<sup>4</sup> | 0.9782 |
| 1.60x10<sup>3</sup> | 3.05 x10<sup>5</sup> | 0.9083 |
| 7.80x10<sup>2</sup> | 1.00 x10<sup>6</sup> | 0.9795 |

### Pseudovirus titration

Pseudovirus titration using ancestral Wuhan spike pseudotyped lentivirus was performed to identify optimal cell density corresponding to high RLU pseudovirus signal. Establishing a high pseudovirus RLU signal is required to create a sufficient signal above the cell-only background of at least 1000-fold in order to determine reportable IC_50_ values that meet the acceptance criteria (19). Pseudovirus was initially diluted 100-fold followed by an 8-step 2-fold serial dilution. A decrease in pseudovirus RLU signal was shown at cell densities above 2.5×10^4^ cells/well and below 1.3×10^4^ cells/well. High RLU values were observed at 1.30×10^4^ cells/well, showing a linear response from the serial dilutions, which were used to justify the cell density selection at 1.30×10^4^ cells/well (Figure 1).

### Clinical Specificity and Sensitivity

Specificity and sensitivity were examined against the ancestral Wuhan and B.1.1.7 spike pseudotyped lentiviruses, in comparison to the gold-standard PRNT assay. Sixty positive samples for SARS-CoV-2 from the NML COVID-19 National Panel and sixty SARS-CoV-2 negative pre-COVID-19 samples were used for the experiment. The results for both parameters against ancestral Wuhan spike pseudotyped lentivirus were acceptable, achieving 100% (95% CI: 94-100) specificity and 100% (95% CI: 94-100) sensitivity. For B.1.1.7 spike pseudotyped lentivirus, sensitivity of 88.3% (95% CI: 77.8 to 94.2) and specificity of 100% (95% CI: 94-100) were achieved (Table 2). A perfect interrater agreement with the PRNT_50_ was demonstrated against the ancestral Wuhan spike pseudotyped lentivirus and almost perfect agreement (κ-value 0.883) shown with the B.1.1.7 spike pseudotyped lentivirus against the PRNT_50_ respectively (Table 2).

**Table 2.**
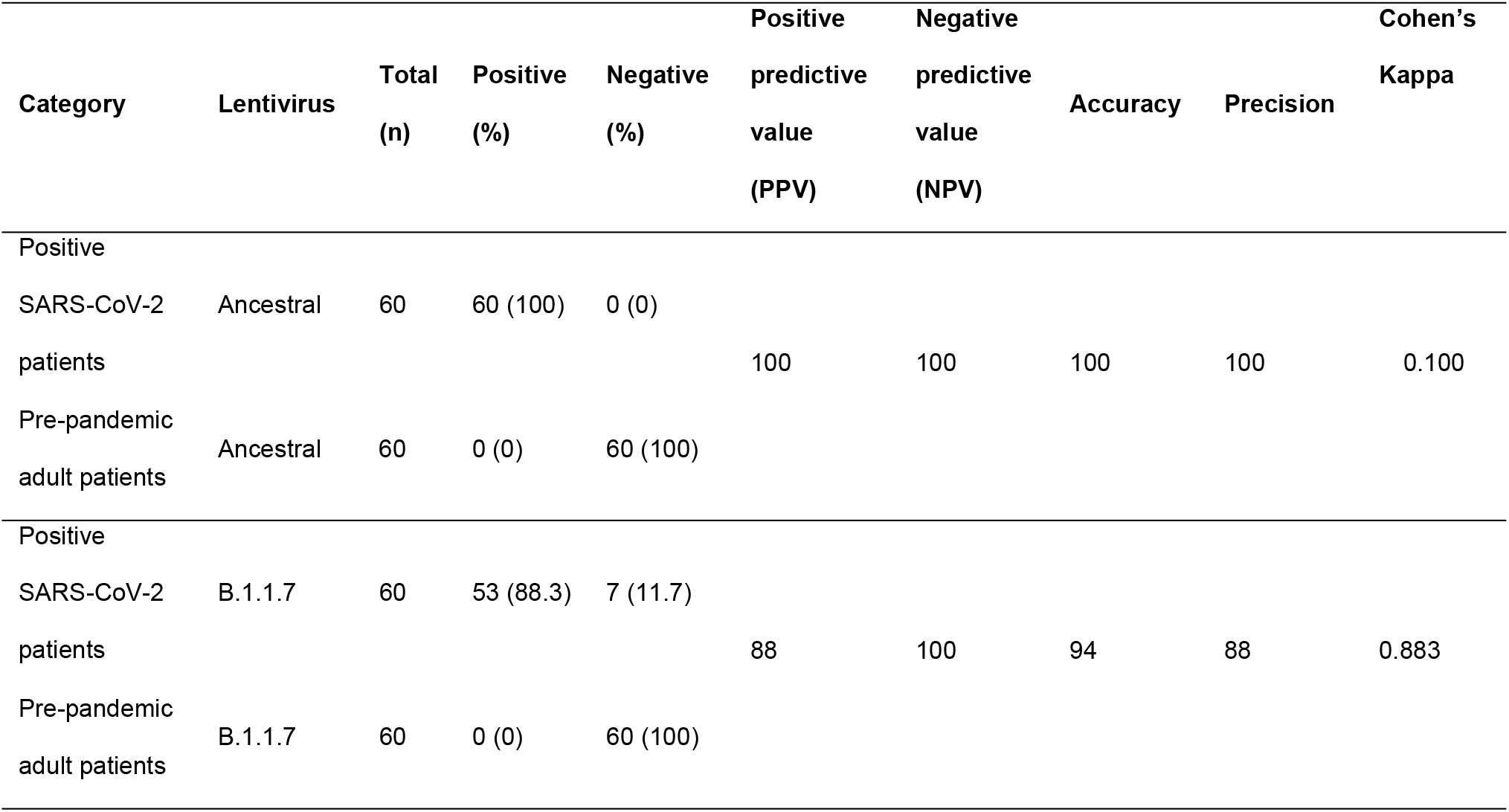
Clinical specificity and sensitivity results based on the comparison between the SCLSNA and the PRNT_50_.

### Validation of the SCLSNA

Guidelines used for the validation were based on the WHO (23) and FDA (27). Validation parameters assessed in this study include precision (repeatability, intermediate precision), robustness, linearity, limit of detection and quantification, as described below. Accuracy was not assessed due to the limitation in accurately comparing reportable values between the IC_50_ of the SCLSNA and PRNT_50_.

#### Precision

##### Repeatability (intra-assay precision)

Repeatability was measured using three concentrations that was based on our in-house PRNT_50_ titer results. The concentrations consisted of “High” (1:≥ 640 PRNT_50_), “Mid” (1:160 PRNT_50_) and “Low” (1:40 PRNT_50_) samples. Analysts processed each sample in triplicate on three separate weeks for a total of nine determinations each (Figure 2). %CV for each concentration were within the acceptance criteria of ≤ 20% CV, with values of 14.21 %CV (High), 12.47% (Mid) and 13.28% CV (Low). Weekly comparisons were within the acceptable range with %CV of 14.44% (Week 1), 18.79% (Week 2) and 9.696% (Week 3).

**Figure 2.**
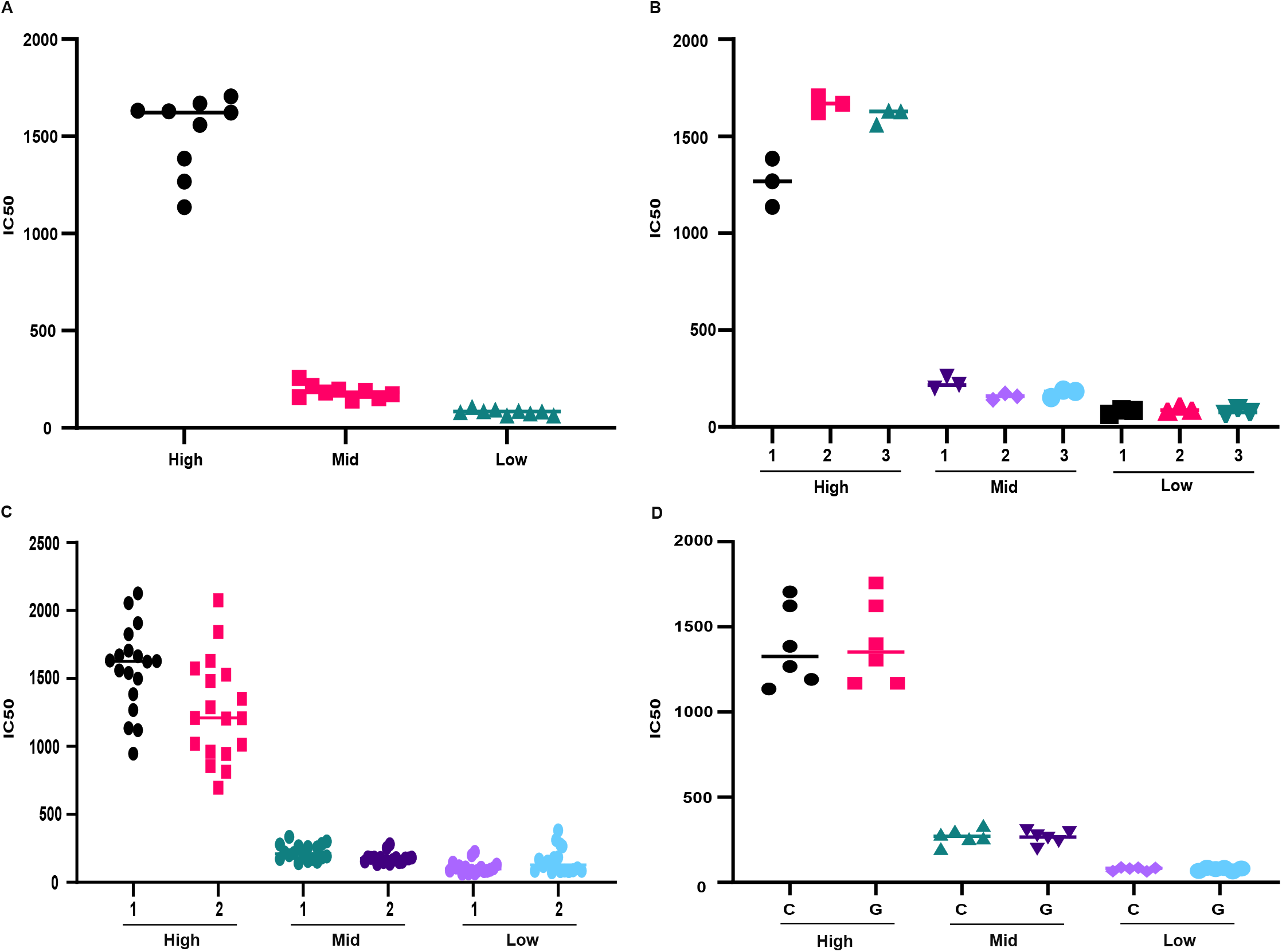
(**A and B)** Intra-assay variability of SCLSNA for one analyst. One analyst tested High (1:≥ 640 PRNT_50_), Mid (1:160 PRNT_50_) and Low (1:40 PRNT_50_) samples from the NML CNP against ancestral Wuhan Spike pseudotyped lentivirus. A) Analyst 1 tested High (1:≥ 640 PRNT_50_), Mid (1:160 PRNT_50_) and Low (1:40 PRNT_50_) samples from the NML CNP against ancestral Wuhan spike pseudotyped lentivirus in triplicate on three separate weeks for nine determinations of each concentration. IC_50_ values were compared and evaluated based on %CV. **(B)** Analyst 1 week to week comparison measuring intra-assay variability using the same High (1:≥ 640 PRNT_50_), Mid (1:160 PRNT_50_) and Low (1:40 PRNT_50_) samples from the NML CNP against ancestral Wuhan Spike pseudotyped lentivirus along with the same conditions and equipment each week. **(C)** Inter-assay variability of SCLSNA between two analysts. Two analysts tested High (1:≥ 640 PRNT_50_), Mid (1:160 PRNT_50_) and Low (1:40 PRNT_50_) samples from the NML CNP against ancestral Wuhan Spike pseudotyped lentivirus. Samples were tested in six replicates on three separate weeks for a total of eighteen determinations each. The solid line represents the mean IC_50_ titer. Results were reported as IC_50_ titers and %CV comparison between analysts were done using GraphPad Prism v.9.3 software. **(D)** Inter-assay variability comparison of the IC_50_ between the Agilent BioTek Cytation 1 (C) and Promega’s GloMax® Navigator microplate luminometer (G). Analyst 1 tested High (1:≥ 640 PRNT_50_), Mid (1:160 PRNT_50_) and Low (1:40 PRNT_50_) samples from the NML CNP against ancestral Wuhan Spike pseudotyped lentivirus. Six replicates were used with each device for eighteen determinations. The solid line represents the mean IC_50_ titer. Results were reported as IC_50_ titers and %CV comparison between devices were done using GraphPad Prism v.9.3 software.

#### Intermediate precision (inter-assay variability)

Inter-assay variability was assessed by comparing the same homogeneous sample between different analysts tested on different weeks. Each analyst tested six replicates of High, Mid and Low samples from the NML CNP on three separate weeks for eighteen determinations (Figure 2, 3). %CV for High and Mid samples between analysts were within acceptable range, with %CV of 15.52% (High) and 16.09% (Mid). %CV for Low did not meet acceptance criteria, with a %CV of 26.42%, indicating slightly higher variation within the Low samples between analysts.

**Figure 3.**
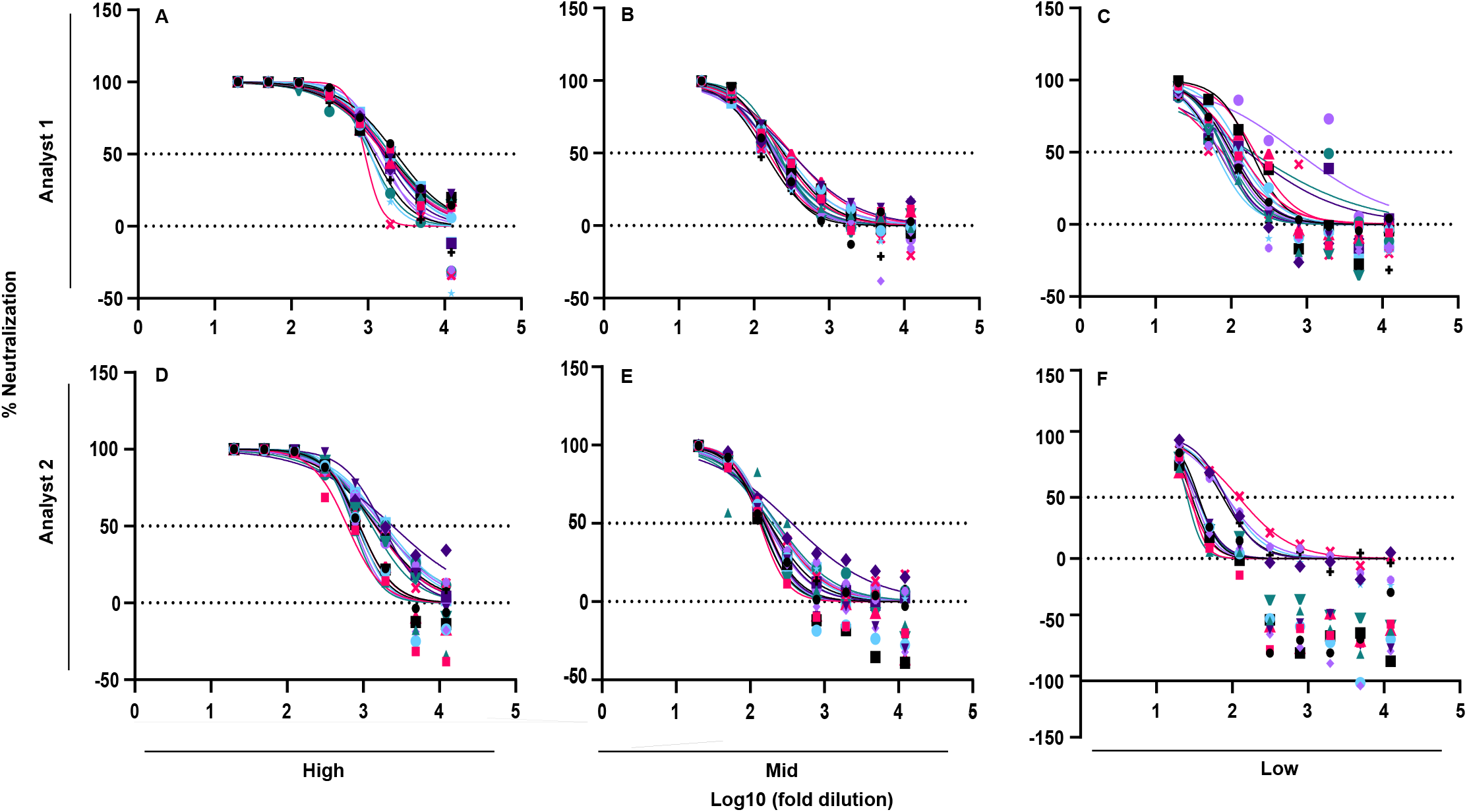
Inter-assay variability of SCLSNA between two analysts. Percent neutralization comparison between two analysts using **(A, D)** High (1:≥ 640 PRNT_50_), **(B, E)** Mid (1:160 PRNT_50_) and **(C, F)** Low (1:40 PRNT_50_) samples from the NML CNP against ancestral Wuhan Spike pseudotyped lentivirus. Samples were tested in six replicates on three separate weeks for a total of fifty-four determinations. IC_50_ titers were determined using GraphPad Prism v.9.3 software.

#### Robustness

Robustness of the SCLSNA was examined to measure the ability of the procedure to provide analytical results of acceptable accuracy and precision under a variety of conditions. In this experiment, a High, Mid and Low sample were tested in triplicate for two independent runs and the detection method was compared using an Agilent BioTek Cytation 1 cell imaging multimode reader device and Promega’s GloMax® Navigator microplate luminometer (Figure 2). Results from the different devices were compared to determine if changes in operational conditions influenced results. Intra-assay variability for each device were below 20% CV and acceptable (Table 3). Inter-assay variability between the devices for each concentration were below 20% CV and passed the criteria (Table 3).

**Table 3.**
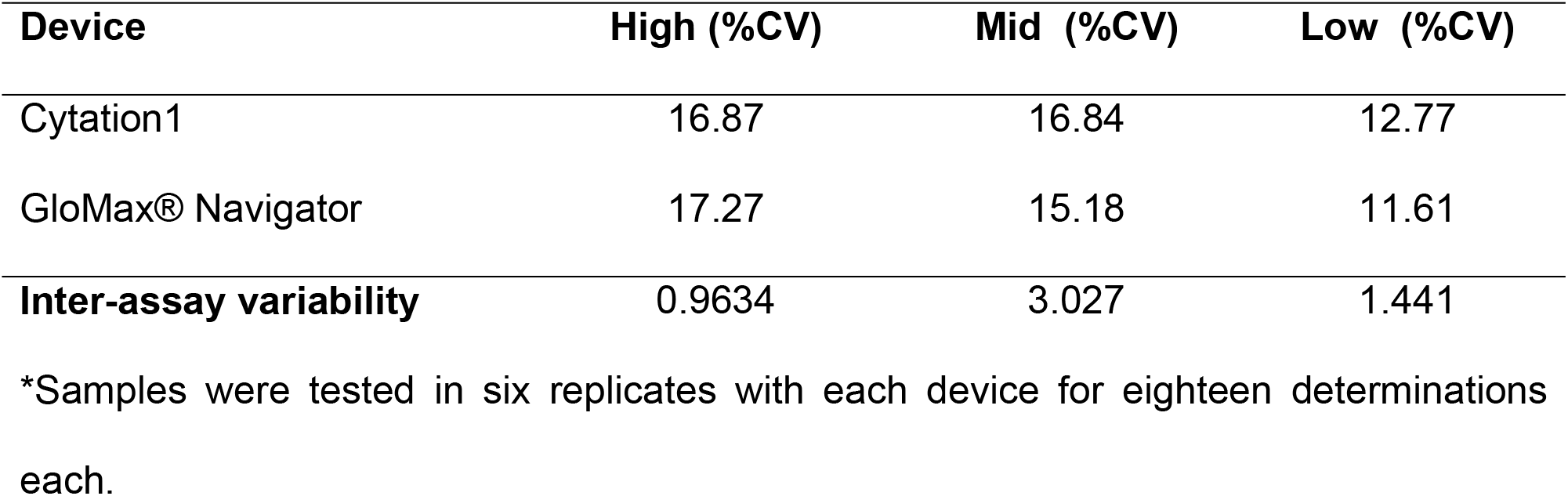
Intra-assay variability analysis and inter-assay variability using the Agilent BioTek Cytation 1 and Promega’s GloMax® Navigator microplate luminometer using a High, Mid and Low sample.

| Device | High (%CV) | Mid (%CV) | Low (%CV) |
| --- | --- | --- | --- |
| Cytation1 | 16.87 | 16.84 | 12.77 |
| GloMax® Navigator | 17.27 | 15.18 | 11.61 |
| **Inter-assay variability** | 0.9634 | 3.027 | 1.441 |

\*Samples were tested in six replicates with each device for eighteen determinations each.

#### Linearity

Linearity was assessed using a WHO international reference panel for anti-SARS-CoV-2 immunoglobulin. The WHO panel consisted of five pooled human plasma samples of High (20/150), Mid (20/148), Low 1 (20/144), Low 2 (20/140) and pre-COVID-19 (20/142) samples. IC_50_ titers obtained from the SCLSNA indicate that they are directly proportional to antibody titers of the WHO reference panel, with IC_50_ values of 331.2 (High), 171.7 (Mid), 107.8 (Low 1), 32.01 (Low 2) and 10 (Pre-COVID-19) (Figure 4 – A, B). SCLSNA IC_50_ values and antibody titers (IU/mL) from the WHO reference panel were compared using Pearson’s correlation coefficient analysis. A strong correlation between the WHO reference panel antibody titers and SCLSNA was observed, with a correlation of r = 0.9210, *p* = 0.0263 (Figure 4C). Linearity was also assessed with pseudovirus addition and RLU. Here, we showed dilutional linearity with the pseudovirus that is observed with decreasing RLUs as the dilution increases (Figure 4D).

**Figure 4.**
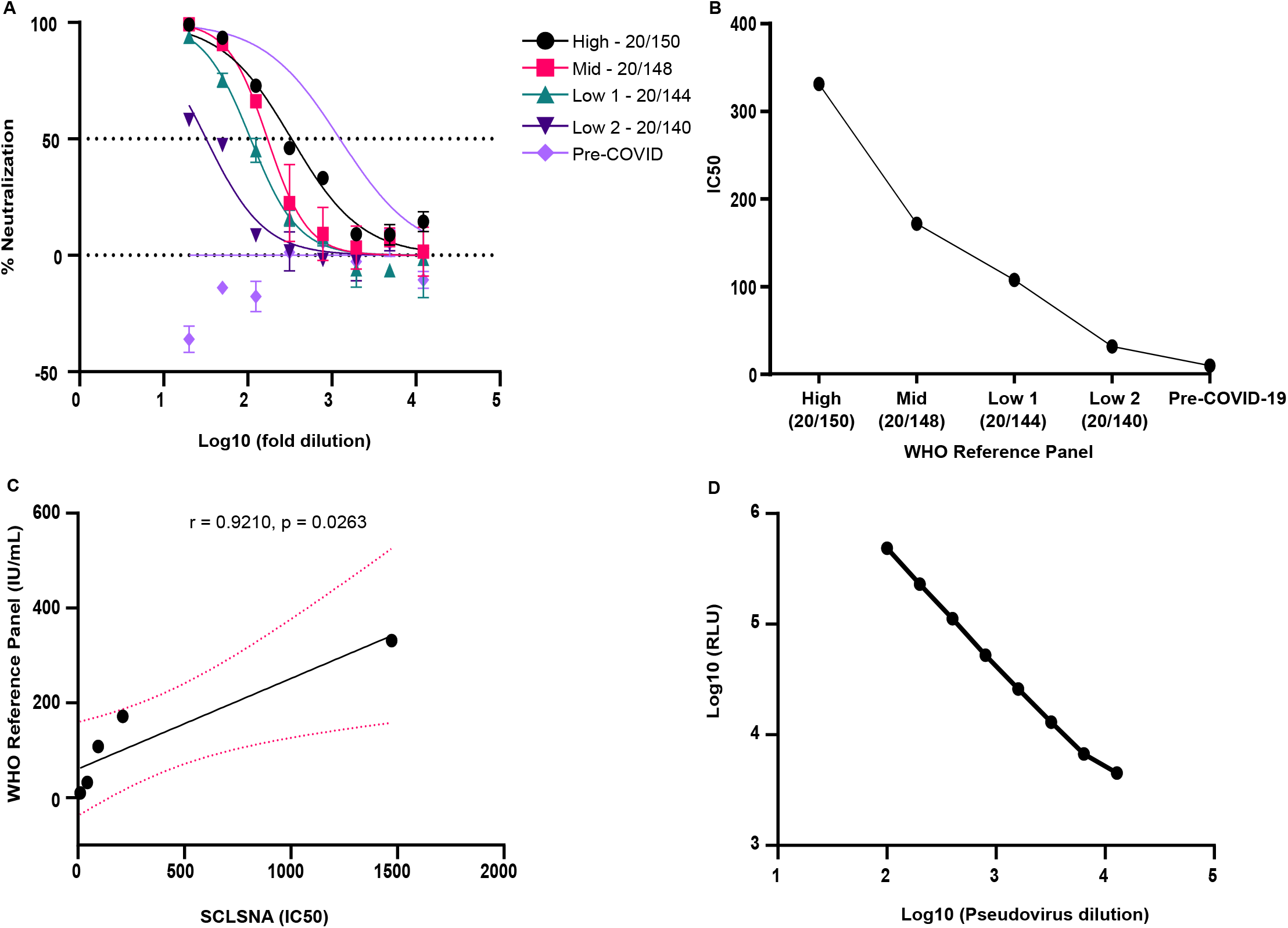
Linearity analysis of the SCLSNA. **(A)** SCLSNA analysis of High, Mid, Low and pre-COVID-19 samples. IC_50_ titers were determined using GraphPad Prism v.9.3. **(B)** Linearity analysis was performed to compare the IC_50_ from the SCLSNA to the antibody titers from the WHO reference panel. **(C)** Correlation analysis between the SCLSNA and the WHO reference panel (IU/mL). **(D)** Linearity analysis of pseudovirus dilution and relative light units. Five reference standards were tested (High, Mid, Low 1, Low 2 and Pre-COVID-19). The solid line indicates the regression line and the dashed line indicates the 95% confidence interval (95% CI: 0.2066 to 0.9949). Pearson’s correlation coefficient and *P*-value are indicated.

**Figure 5.**
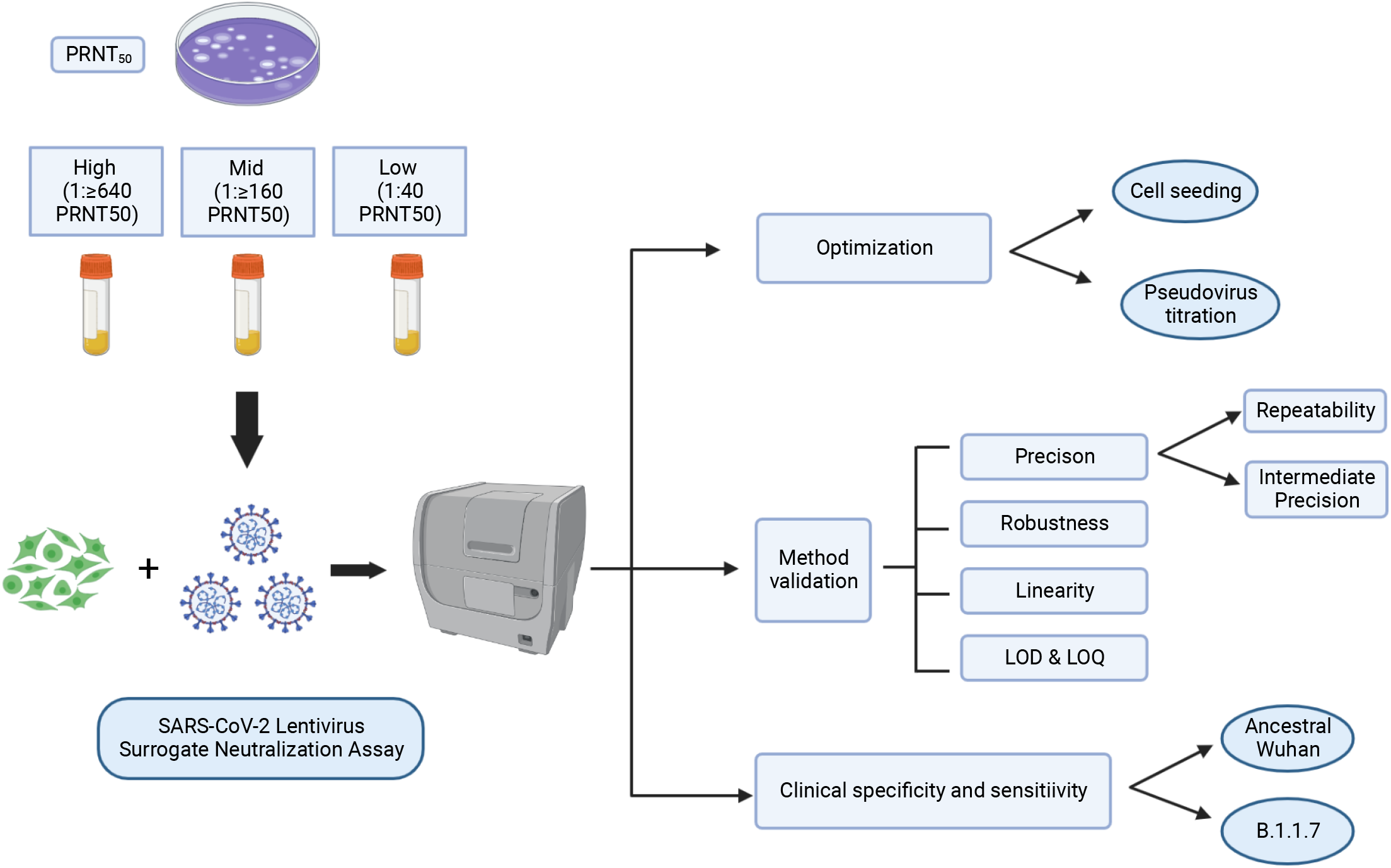
Flow chart schematic of the SCLSNA validation study design. High (1:≥ 640 PRNT_50_), Mid (1:160 PRNT_50_) and Low (1:40 PRNT_50_) samples from the NML CNP were tested against ancestral Wuhan Spike pseudotyped lentivirus by the SCLSNA. Assay optimization was conducted to determine optimal HEK293T/ACE2-TMPRSS2 cell seeding density and a pseudovirus titration was performed to confirm optimal cell seeding density. All High, Mid and Low samples were used for the method validation to test precision, robustness, linearity, LOD and LOQ. Direct comparison of the SCLSNA to the gold-standard PRNT was determined through the clinical specificity and sensitivity using an ancestral Wuhan and B.1.1.7 Spike pseudotyped lentivirus. Created with BioRender.com.

#### Limit of Detection and Limit of Quantification

LOD and LOQ for the SCLSNA was determined using thirty-six samples from the NML CNP that were pre-COVID-19 and negative for SARS-CoV-2 (as verified by the PRNT). The standard deviation determined from the mean IC_50_ values of the negative samples resulted in a LOD of 19.60 and a LOQ of 65.32 which were three and ten times the standard deviation respectively. We used a cut-off of < 20 for negative samples and assigned them a nominal value of 10. This was done to distinguish negative from positive results in our qualitative representation of our results.

## Discussion

We have shown the SCLSNA to be a suitable alternative to the gold-standard PRNT. In this validation study, we established acceptable validation parameters for precision, robustness and linearity while optimizing and displaying sensitivity and specificity with the SCLSNA that were comparable to the PRNT. Other studies have previously validated similar versions of surrogate neutralization assays but the goal of this study was to expand the validation parameters tested and include a sample concentration range based off of the PRNT to further confirm reliability and strength of the SCLSNA as a comparative approach to the PRNT (9, 10, 17).

Overall, good precision was shown throughout the validation study. Previous studies have also shown good precision for both intra-assay and inter-assay variability (9, 10, 28, 29), but one key difference in our approach was the use of a broad concentration range of samples. Incorporation of High (1:≥ 640 PRNT_50_), Mid (1:160 PRNT_50_) and Low (1:40 PRNT_50_) samples allowed us to directly compare samples between the SCLSNA and PRNT, allowing for a thorough analysis on precision within and between analysts. Neerukonda S. *et*.*al*., 2021, used a similar broad-based approach on sample concentrations; however, more variation was detected in their intermediate precision, which was greater than what was shown in our study (17). We also detected higher than expected variation within the Low samples between analysts, which may be due to the specificity of binding inherent within the SCLSNA, which focuses solely on the receptor binding domain and spike regions of SARS-CoV-2, as opposed to non-specific binding to live SARS-CoV-2 that may be found in plasma samples at lower dilutions (30, 31). Spike protein density within a pseudotyped lentivirus may also be different than live SARS-CoV-2, which may result in a decreased amount of neutralization, particularly in low titer samples (32). Despite the outcome observed in the Low sample comparison, we were still able to show acceptable precision for intra- and inter-assay variability in each concentration range, along with low variation within and between analysts.

Another strategy we employed in our validation was the use of multiple luminescence detectors to achieve robustness; we were able to achieve acceptable robustness from the low variation seen across different luminescence detectors. To our knowledge, this approach has not been evaluated in previous studies and herein we showed low variation across several different devices, indicating the flexibility of the SCLSNA in its performance and capabilities. One limitation in our robustness analysis was the inability to compare assay performance in different laboratories due to logistical challenges and unavailability at the time of the study. We opted to test robustness using different luminescence detectors as an alternative and to confirm the reliability of the assay.

Linearity of the SCLSNA was demonstrated after comparison with the WHO international reference panel. The SCLSNA closely approximated the expected concentrations of the standard showing a strong correlation between the SCLSNA and NIBSC reference standard. A study conducted by Yu J. *et*.*al*., 2021, achieved dilutional linearity which was also confirmed in our study, highlighting the ability of the SCLSNA in measuring expected values that are directly proportional to the amount of pseudovirus used (29).

Optimization of the SCLSNA established the optimal seeding density as previously shown in other studies (9, 10). In those studies, Huh7 and BHK21-hACE2 cell lines were used, in contrast to our studies where we opted to use HEK293T/ACE2-TMPRSS2 cells due to the ability of TMPRSS2 to prime spike proteins on pseudotyped lentiviruses, likely increasing infectivity (33). Another important component to the HEK293T/ACE2-TMPRSS2 cells was the addition of polybrene, which is a polycationic agent that helps facilitate pseudovirus cell entry (34). In our study, we optimized the SCLSNA to help establish consistency, minimize variation and ensure our method was performing at optimal levels.

To determine if the SCLSNA can perform similarly to the gold-standard PRNT, we conducted a comparability study of the SCLSNA to the PRNT through analysis of clinical sensitivity and specificity. Using either ancestral Wuhan or B.1.1.7 spike pseudotyped lentivirus, we were able to achieve acceptable sensitivity and specificity. We observed very high sensitivity and specificity for ancestral Wuhan spike pseudotyped lentivirus; however, we also saw a decrease in sensitivity with the B.1.1.7 spike pseudotyped lentivirus. This may be attributed to a reduced level of neutralization found against the B.1.1.7 variant, as evidenced by reduced neutralization activities of various monoclonal antibodies (35, 36). In addition, convalescent sera and vaccine-induced antibody responses are still effective against the B.1.1.7 variant but the immune response may vary in comparison to the ancestral Wuhan pseudotyped lentivirus (35, 36). To determine correlation between the SCLSNA and PRNT using SARS-CoV-2 convalescent patient samples, we conducted a correlation assessment, but the correlation coefficient was low (data not shown). The NML COVID-19 National Panel sample set used in this study consisted mainly of one antibody titer range at lower neutralization titers (1:80 PRNT_50_) from convalescent donors who were naturally infected with SARS-CoV-2 before the vaccine was available, making it difficult to achieve a proper correlation analysis. As a result, we used clinical sensitivity and specificity as a measure for comparison and were able to show good comparability to the PRNT, as seen in other studies (12, 28).

This validation study successfully achieved acceptable criteria in all the parameters tested, proving the SCLSNA to be a reliable pre-screening approach to the PRNT. One of the key advantages to the SCLSNA is the use of a SARS-CoV-2 pseudotyped lentivirus generation platform. The lentivirus system enables an efficient and quick TAT for generating lentiviruses pseudotyped with the target-of-interest. This is particularly important during the pandemic with the emergence of novel variants of concern to which pre-existing NAbs may be less effective. As well, all plasmids used in generating our pseudotyped lentiviruses are commercially available, making for a convenient and time-saving approach in comparison to custom designed plasmids which are more time consuming to prepare (15). In addition, the in-house generation of pseudotyped lentivirus is a faster approach than the generation of live virus in a BSL-3 setting, which may take time to successfully rescue live virus and optimize assay conditions.

Another key advantage is the quantitative output of the SCLSNA. The data generated from this assay gives a more precise antibody titer with the IC_50_ rather than a visual determination of antibody titer by the PRNT method, which is considered more subjective as technical staff must be carefully trained to accurately identify plaque formation by visual means (28). It is also difficult to obtain an end-point dilution for a large sample set, especially ones containing high antibody titers, resulting in a broad estimation of antibody titer and a “cut-off” value assigned in the reported data affecting the overall precision of the results. Data generated by the PRNT can be more subjective across different analysts, further decreasing accuracy and consistency of results (8). Furthermore, sample throughput for the PRNT is limited to processing a lower amount of samples which requires plates with larger well sizes and manual labor (16, 37). The plaque morphology with each new variant changes resulting in subjectivity amongst analysts and inaccurate reporting of results (38, 39). The focus reduction neutralization test (FRNT) has been used as an alternative to the PRNT but limitations such as the need for a BSL-3 facility and qualitative analysis still remain (40–42).In contrast, the SCLSNA can be used for high-throughput, automated sample processing in 96 to 384 well plate formats (16, 37).

Validation of the SCLSNA provides an alternative neutralizing antibody platform to support or potentially replace the PRNT gold-standard method. The SCLSNA does not require the handling of live SARS-CoV-2 virus in a BSL-3 facility, providing for a safer work environment, is less tedious and has a faster TAT for sample processing to reporting of results. The quantitative analysis that is achievable by the SCLSNA increases its precision, making it a reliable approach to the limitations found inherent within the PRNT. The validation parameters tested in this study met the previously established acceptance criteria, making the SCLSNA a suitable alternative to the PRNT.

## References

1. Thompson MG, Stenehjem E, Grannis S, Ball SW, Naleway AL, Ong TC, DeSilva MB, Natarajan K, Bozio CH, Lewis N, Dascomb K, Dixon BE, Birch RJ, Irving SA, Rao S, Kharbanda E, Han J, Reynolds S, Goddard K, Grisel N, Fadel WF, Levy ME, Ferdinands J, Fireman B, Arndorfer J, Valvi NR, Rowley EA, Patel P, Zerbo O, Griggs EP, Porter RM, Demarco M, Blanton L, Steffens A, Zhuang Y, Olson N, Barron M, Shifflett P, Schrag SJ, Verani JR, Fry A, Gaglani M, Azziz-Baumgartner E, Klein NP. 2021. Effectiveness of Covid-19 Vaccines in Ambulatory and Inpatient Care Settings. N Engl J Med 385:1355–1371.

2. Dagan N, Barda N, Kepten E, Miron O, Perchik S, Katz MA, Hernán MA, Lipsitch M, Reis B, Balicer RD. 2021. BNT162b2 mRNA Covid-19 Vaccine in a Nationwide Mass Vaccination Setting. N Engl J Med 384:1412–1423.

3. Tenforde MW, Patel MM, Ginde AA, Douin DJ, Talbot HK, Casey JD, Mohr NM, Zepeski A, Gaglani M, McNeal T, Ghamande S, Shapiro NI, Gibbs KW, Files DC, Hager DN, Shehu A, Prekker ME, Erickson HL, Exline MC, Gong MN, Mohamed A, Henning DJ, Steingrub JS, Peltan ID, Brown SM, Martin ET, Monto AS, Khan A, Hough CL, Busse LW, ten Lohuis CC, Duggal A, Wilson JG, Gordon AJ, Qadir N, Chang SY, Mallow C, Gershengorn HB, Babcock HM, Kwon JH, Halasa N, Chappell JD, Lauring AS, Grijalva CG, Rice TW, Jones ID, Stubblefield WB, Baughman A, Womack KN, Lindsell CJ, Hart KW, Zhu Y, Olson SM, Stephenson M, Schrag SJ, Kobayashi M, Verani JR, Self WH. 2021. Effectiveness of Severe Acute Respiratory Syndrome Coronavirus 2 Messenger RNA Vaccines for Preventing Coronavirus Disease 2019 Hospitalizations in the United States. Clin Infect Dis 2:1–16.

4. Liu Y, Liu J, Shi P-Y. 2022. SARS-CoV-2 Variants and Vaccination. Zoonoses 2:1–9.

5. Harvey WT, Carabelli AM, Jackson B, Gupta RK, Thomson EC, Harrison EM, Ludden C, Reeve R, Rambaut A, Peacock SJ, Robertson DL. 2021. SARS-CoV-2 variants, spike mutations and immune escape. Nat Rev Microbiol 19:409–424.

6. Takeshita M, Nishina N, Moriyama S, Takahashi Y, Ishii M, Saya H, Kondo Y, Kaneko Y, Suzuki K, Fukunaga K, Takeuchi T, Keio Donner Project. 2022. Immune evasion and chronological decrease in titer of neutralizing antibody against SARS-CoV-2 and its variants of concerns in COVID-19 patients. Clin Immunol 108999.

7. Wang W, Kaelber DC, Xu R, Berger NA. 2022. Breakthrough SARS-CoV-2 Infections, Hospitalizations, and Mortality in Vaccinated Patients With Cancer in the US Between December 2020 and November 2021. JAMA Oncol https://doi.org/10.1001/jamaoncol.2022.1096.

8. Valcourt EJ, Manguiat K, Robinson A, Lin Y-C, Abe KT, Mubareka S, Shigayeva A, Zhong Z, Girardin RC, DuPuis A, Payne A, McDonough K, Wang Z, Gasser R, Laumaea A, Benlarbi M, Richard J, Prévost J, Anand SP, Dimitrova K, Phillipson C, Evans DH, McGeer A, Gingras A-C, Liang C, Petric M, Sekirov I, Morshed M, Finzi A, Drebot M, Wood H. 2021. Evaluating Humoral Immunity against SARS-CoV-2: Validation of a Plaque-Reduction Neutralization Test and a Multilaboratory Comparison of Conventional and Surrogate Neutralization Assays. Microbiol Spectr 9:1–15.

9. Nie J, Li Q, Wu J, Zhao C, Hao H, Liu H, Zhang L, Nie L, Qin H, Wang M, Lu Q, Li X, Sun Q, Liu J, Fan C, Huang W, Xu M, Wang Y. 2020. Establishment and validation of a pseudovirus neutralization assay for SARS-CoV-2. Emerg Microbes Infect https://doi.org/10.1080/22221751.2020.1743767.

10. Xiong HL, Wu YT, Cao JL, Yang R, Liu YX, Ma J, Qiao XY, Yao XY, Zhang BH, Zhang YL, Hou WH, Shi Y, Xu JJ, Zhang L, Wang SJ, Fu BR, Yang T, Ge SX, Zhang J, Yuan Q, Huang BY, Li ZY, Zhang TY, Xia NS. 2020. Robust neutralization assay based on SARS-CoV-2 S-protein-bearing vesicular stomatitis virus (VSV) pseudovirus and ACE2-overexpressing BHK21 cells. Emerg Microbes Infect 9:2105–2113.

11. Meyer B, Reimerink J, Torriani G, Brouwer F, Godeke GJ, Yerly S, Hoogerwerf M, Vuilleumier N, Kaiser L, Eckerle I, Reusken C. 2020. Validation and clinical evaluation of a SARS-CoV-2 surrogate virus neutralisation test (sVNT). Emerg Microbes Infect 9:2394–2403.

12. Yang R, Huang B, A R, Li W, Wang W, Deng Y, Tan W. 2020. Development and effectiveness of pseudotyped SARS-CoV-2 system as determined by neutralizing efficiency and entry inhibition test in vitro. Biosaf Heal 2:226–231.

13. Sekirov I, Petric M, Carruthers E, Lawrence D, Pidduck T, Kustra J, Laley J, Lee M-K, Chahil N, Mak A, Levett PN, Mendoza E, Wood H, Drebot M, Krajden M, Morshed M. 2021. Performance comparison of micro-neutralization assays based on surrogate SARS-CoV-2 and WT SARS-CoV-2 in assessing virus-neutralizing capacity of anti-SARS-CoV-2 antibodies. Access Microbiol 3:1–4.

14. Maeda A, Maeda J. 2013. Review of diagnostic plaque reduction neutralization tests for flavivirus infection. Vet J 195:33–40.

15. Wang S, Liu L, Wang C, Wang Z, Duan X, Chen G, Zhou H, Shao H. 2022. Establishment of a pseudovirus neutralization assay based on SARS-CoV-2 S protein incorporated into lentiviral particles. Biosaf Heal 4:38–44.

16. Sholukh AM, Fiore-Gartland A, Ford ES, Miner MD, Hou YJ, Tse L V., Kaiser H, Zhu H, Lu J, Madarampalli B, Park A, Lempp FA, St. Germain R, Bossard EL, Kee JJ, Diem K, Stuart AB, Rupert PB, Brock C, Buerger M, Doll MK, Randhawa AK, Stamatatos L, Strong RK, McLaughlin C, Huang ML, Jerome KR, Baric RS, Montefiori D, Corey L. 2021. Evaluation of cell-based and surrogate SARS-CoV-2 neutralization assays. J Clin Microbiol 59.

17. Neerukonda SN, Vassell R, Herrup R, Liu S, Wang T, Takeda K, Yang Y, Lin TL, Wang W, Weiss CD. 2021. Establishment of a well-characterized SARSCoV-2 lentiviral pseudovirus neutralization assay using 293T cells with stable expression of ACE2 and TMPRSS2. PLoS One 16:1–19.

18. Tan CW, Chia WN, Qin X, Liu P, Chen MIC, Tiu C, Hu Z, Chen VCW, Young BE, Sia WR, Tan YJ, Foo R, Yi Y, Lye DC, Anderson DE, Wang LF. 2020. A SARS-CoV-2 surrogate virus neutralization test based on antibody-mediated blockage of ACE2–spike protein–protein interaction. Nat Biotechnol 38:1073–1078.

19. Crawford KHD, Eguia R, Dingens AS, Loes AN, Malone KD, Wolf CR, Chu HY, Tortorici MA, Veesler D, Murphy M, Pettie D, King NP, Balazs AB, Bloom JD. 2020. Protocol and reagents for pseudotyping lentiviral particles with SARS-CoV-2 spike protein for neutralization assays. Viruses 12.

20. Ferrara F, Temperton N. 2018. Pseudotype neutralization assays: From laboratory bench to data analysis. Methods Protoc https://doi.org/10.3390/mps1010008.

21. Schlimgen R, Howard J, Wooley D, Thompson M, Baden LR, Yang OO, Christiani DC, Mostoslavsky G, Diamond D V., Duane EG, Byers K, Winters T, Gelfand JA, Fujimoto G, Hudson TW, Vyas JM. 2016. Risks associated with lentiviral vector exposures and prevention strategies. J Occup Environ Med 58:1159–1166.

22. Imre S, Vlase L, Muntean DL. 2008. Bioanalytical method validation. Rev Rom Med Lab 10:13–21.

23. World Health Organization. 2018. World Health Organization (WHO). Guidelines on validation - Appendix 4 - Analytical method validation. 1–11.

24. Drews SJ, Devine D V., McManus J, Mendoza E, Manguiat K, Wood H, Girardin R, Dupuis A, McDonough K, Drebot M. 2021. A trend of dropping anti-SARS-CoV-2 plaque reduction neutralization test titers over time in Canadian convalescent plasma donors. Transfusion 61:1440–1446.

25. Ferrara F, Temperton N. 2018. Pseudotype neutralization assays: From laboratory bench to data analysis. Methods Protoc 1:1–16.

26. Wang S, Sakhatskyy P, Chou THW, Lu S. 2005. Assays for the assessment of neutralizing antibody activities against Severe Acute Respiratory Syndrome (SARS) associated coronavirus (SCV). J Immunol Methods 301:21–30.

27. U.S. Department of Health and Human Services, Food and Drug Administration. 2018. Bioanalytical method validation guidance for industry. US Dep Heal Hum Serv Food Drug Adm 1–41.

28. Tolah AMK, Sohrab SS, Tolah KMK, Hassan AM, El-Kafrawy SA, Azhar EI. 2021. Evaluation of a pseudovirus neutralization assay for sars-cov-2 and correlation with live virus-based micro neutralization assay. Diagnostics 11.

29. Yu J, Li Z, He X, Gebre MS, Bondzie EA, Wan H, Jacob-dolan C. 2021. Deletion of the SARS-CoV-2 Spike Cytoplasmic Tail Increases.

30. Sarzotti-Kelsoe M, Bailer RT, Turk E, Lin C li, Bilska M, Greene KM, Gao H, Todd CA, Ozaki DA, Seaman MS, Mascola JR, Montefiori DC. 2014. Optimization and validation of the TZM-bl assay for standardized assessments of neutralizing antibodies against HIV-1. J Immunol Methods 409:131–146.

31. Brian Hetrick, Linda D. Chilin, Sijia He, Deemah Dabbagh, Farhang Alem, Aarthi Narayanan, Alessandra Luchini, Tuanjie Li, Xuefeng Liu JC. 2022. Development of a hybrid alphavirus-SARS-CoV-2 pseudovirion for rapid quantification of neutralization antibodies and antiviral drugs. Cell Reports Methods 19–21.

32. Schmidt F, Weisblum Y, Muecksch F, Hoffmann HH, Michailidis E, Lorenzi JCC, Mendoza P, Rutkowska M, Bednarski E, Gaebler C, Agudelo M, Cho A, Wang Z, Gazumyan A, Cipolla M, Caskey M, Robbiani DF, Nussenzweig MC, Rice CM, Hatziioannou T, Bieniasz PD. 2020. Measuring SARS-CoV-2 neutralizing antibody activity using pseudotyped and chimeric viruses. J Exp Med 217.

33. Hoffmann M, Kleine-Weber H, Schroeder S, Krüger N, Herrler T, Erichsen S, Schiergens TS, Herrler G, Wu NH, Nitsche A, Müller MA, Drosten C, Pöhlmann S. 2020. SARS-CoV-2 Cell Entry Depends on ACE2 and TMPRSS2 and Is Blocked by a Clinically Proven Protease Inhibitor. Cell 181:271-280.e8.

34. Denning W, Das S, Guo S, Xu J, Kappes JC, Hel Z. 2013. Optimization of the transductional efficiency of lentiviral vectors: effect of sera and polycations. Mol Biotechnol 53:308–314.

35. Shen X, Tang H, McDanal C, Wagh K, Fischer W, Theiler J, Yoon H, Li D, Haynes BF, Sanders KO, Gnanakaran S, Hengartner N, Pajon R, Smith G, Glenn GM, Korber B, Montefiori DC. 2021. SARS-CoV-2 variant B.1.1.7 is susceptible to neutralizing antibodies elicited by ancestral spike vaccines. Cell Host Microbe 29:529-539.e3.

36. Muik A, Wallisch AK, Sänger B, Swanson KA, Mühl J, Chen W, Cai H, Maurus D, Sarkar R, Türeci Ö, Dormitzer PR, Şahin U. 2021. Neutralization of SARS-CoV-2 lineage B.1.1.7 pseudovirus by BNT162b2 vaccine–elicited human sera. Science (80-) 371:1152–1153.

37. Sarzotti-Kelsoe M, Cox J, Cleland N, Denny T, Hural J, Needham L, Ozaki D, Rodriguez-Chavez IR, Stevens G, Stiles T, Tarragona-Fiol T, Simkins A. 2009. Evaluation and recommendations on good clinical laboratory practice guidelines for phase I-III clinical trials. PLoS Med 6:1–5.

38. Jeong GU, Yoon GY, Moon HW, Lee W, Hwang I, Kim H, Kim K Do, Kim C, Ahn DG, Kim BT, Kim SJ, Kwon YC. 2022. Comparison of plaque size, thermal stability, and replication rate among SARS-CoV-2 variants of concern. Viruses 14.

39. Khandelwal N, Chander Y, Kumar R, Nagori H, Verma A, Mittal P, Riyesh T, Kamboj S, Verma SS, Khatreja S, Pal Y, Gulati BR, Tripathi BN, Barua S, Kumar N. 2021. Studies on Growth Characteristics and Cross-Neutralization of Wild-Type and Delta SARS-CoV-2 From Hisar (India). Front Cell Infect Microbiol 11.

40. K VSbDjLsDaNb. 2010. Development of a focus reduction neutralization test (FRNT) for detection of mumps virus neutralizing antibodies. J Virol Methods 163:153–156.

41. Abigail Vanderheiden 1 2 3, Venkata Viswanadh Edara 1 2 3, Katharine Floyd 1 2 3, Robert C Kauffman 2 4, Grace Mantus 2 4, Evan Anderson 1, Nadine Rouphael 2 5, Sri Edupuganti 2 5, Pei-Yong Shi 6, Vineet D Menachery 7, Jens Wrammert 2 4 MSS 1 2 3. 2020. Development of a Rapid Focus Reduction Neutralization Test Assay for Measuring SARS-CoV-2 Neutralizing Antibodies. Curr Protoc Immunol 131.

42. Guruprasad R Medigeshi 1, Gaurav Batra 1, Deepika Rathna Murugesan 1, Ramachandran Thiruvengadam 1, Souvick Chattopadhyay 1, Bhabatosh Das 1, Mudita Gosain 1, Ayushi 1, Janmejay Singh 1, Anantharaj Anbalagan 1, Heena Shaman 1, Kamal Pargai 1, Farha Mehdi PKG 3. 2022. Sub-optimal neutralisation of omicron (B.1.1.529) variant by antibodies induced by vaccine alone or SARS-CoV-2 Infection plus vaccine (hybrid immunity) post 6-months. EBioMedicine 78.

